# Theta sequences support real-time spatial memory retrieval

**DOI:** 10.64898/2026.10.18.759880

**Authors:** Livia Stein, Guillaume Etter

## Abstract

Oscillations play a critical role in orchestrating the timing of neural activity during cognitive processes. In the hippocampus, place cells encode spatial locations, and their activity is organized by theta oscillations into sequences that represent past, present, and potential future states. Recently, scrambling theta oscillations in vivo was shown to impair memory retrieval during navigation and decision-making, while leaving place cell activity intact. This surprising result highlights that the exact mechanisms underlying real-time memory encoding and retrieval in the hippocampus remain undefined. Here, we use a spiking neural network model of hippocampal theta sequences to evaluate the consequences of theta frequency scrambling on both representational learning and real-time spatial memory retrieval. We first show that frequency scrambling disrupts theta sequences and phase precession. Specifically, increased frequency scrambling was associated with a gradually degraded relationship between theta phase and spike timing, and ultimately abolished the predictive nature of place fields. When hippocampal spikes are required to solve a T-maze alternation task, the number of successful trials and navigation efficiency decreased with increasing values of frequency scrambling. These results could advance our understanding of how theta modulated neural activity supports memory-guided decision-making in real time and demonstrate why navigation fails when disrupting theta, despite intact spatial representations.

## Introduction

Efficient and goal-directed navigation depends on the ability to use learned spatial representations to guide behavior and support future choices[1, 2]. The hippocampus is thought to play a key role in these processes due to its ability to represent allocentric (self-location) information through place cells - pyramidal neurons that fire selectively when the animal occupies a particular location [3]. At the population level, the location-specific firing patterns (place fields) of place cells are presumed to form a spatial cognitive map of the environment that supports orientation and flexible navigation [4, 5]. In addition to a rate code, which reflects the animal’s position through the cell’s firing rate [4, 6], hippocampal representations also rely on a temporal code, which critically depends on theta oscillations [6–8], a rhythmic pattern in the 6 to 12 Hz frequency band that is prominent in the hippocampal local field potential (LFP) [9, 10], and which is paced by upstream medial septal inputs [10–12]. At the level of individual theta cycles, theta oscillations organize place cell firing into sequences that span past, present and potential future locations along the animal’s trajectory [13–15] and provide an online representation of the animal’s experience. One mechanism thought to underlie this temporal compression is theta phase precession [13, 16, 17], whereby place cells spike at progressively earlier phases of successive theta cycles as the animal traverses their place fields. Beyond reflecting ongoing experience, theta sequences have been linked to goal-directed navigation [18–20], online spatial planning [19], continuous simulation of possible futures [21], and decision points in a maze [22, 23], where they can sweep ahead of the animal to represent different options in alternating theta cycles before a choice is made [22]. Together, this suggests that theta sequences may serve as a neural format that supports spatial (goal-directed) planning by representing alternative paths during navigation [18–20, 22, 24, 25] and that may support both the encoding and online retrieval of spatial representations. Previous studies have manipulated theta oscillations in vivo through pharmacological inhibition [26, 27], optogenetic inhibition [28, 29] or optogenetic stimulation [30, 31] targeting neurons in the medial septum. Remarkably, such manipulations were found to have an effect on either or both memory encoding [27–29] and retrieval [30] during spatial navigation and decision-making, while leaving spatial representations and firing selectivity of place cells surprisingly intact [30]. In particular, optogenetic frequency scrambling, which preserves theta power while disrupting phase coordination, suggests that theta rhythmicity, rather than theta power alone, is necessary for successful retrieval of spatial information [30]. Complementary computational work further shows that theta oscillations are critical for learning predictive spatial representations, and that without theta the learned spatial map loses its predictive structure [32]. Taken together, these findings and the dissociation between place fields and spatiotemporal codes suggest that spatial rate codes alone are insufficient for maintaining successful memory-guided navigation and that theta-dependent temporal organization of neural activity may be required to transform hippocampal representations into a format that supports online retrieval and effective readout by downstream regions. However, the exact mechanism through which graded theta frequency scrambling alters spike timing, theta sequences, and place cell dependent navigation remain incompletely understood. To address this, we examined how frequency scrambling affects the online retrieval of encoded spatial representations during navigation using a spiking neural network model. By systematically increasing the degree of phase scrambling, we show that spike timing progressively detaches from theta phase and theta sequences degenerate. Behaviorally, this was reflected in reduced navigation stability and efficiency in a T-maze alternation task, visualizing a progressively impaired ability to retrieve and deploy information encoded in spatial representations.

## Results

### Frequency scrambling disrupts theta sequences and spike timing

To investigate the role of theta oscillations in memory retrieval during spatial navigation and decision making, we adapted the hippocampal spiking neural network model developed by George *et al*. [32]. The model consists of one layer of CA3 place cells and a downstream layer of CA1 place cells that are connected via synaptic weights, learned through spike-time-dependent plasticity (STDP) as an agent explores a maze (Fig. 1**a**). During exploration, the location *x*(*t*) of the agent in the maze is encoded by the firing of a population of CA3 place cells, each of which has a thresholded Gaussian receptive field 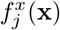 with a set radius and peak firing rate. The model further encompasses a global theta oscillation by which each place cell’s firing rate is modulated. Specifically, a phase precession factor 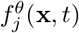 suppresses individual place cell firing for all but a preferred phase during consecutive theta cycles and this preferred phase shifts to progressively earlier phases as the agent traverses the cell’s receptive field. At the population level, the restriction of spikes to phases reflecting the position of the agent relative to the location encoded by the cell results in theta sequences. In other words, spikes from different cells along the agent’s path are sequentially ordered in time by phase in individual cycles, effectively giving rise to sequences (Fig. 1**b**).

**Figure 1:**
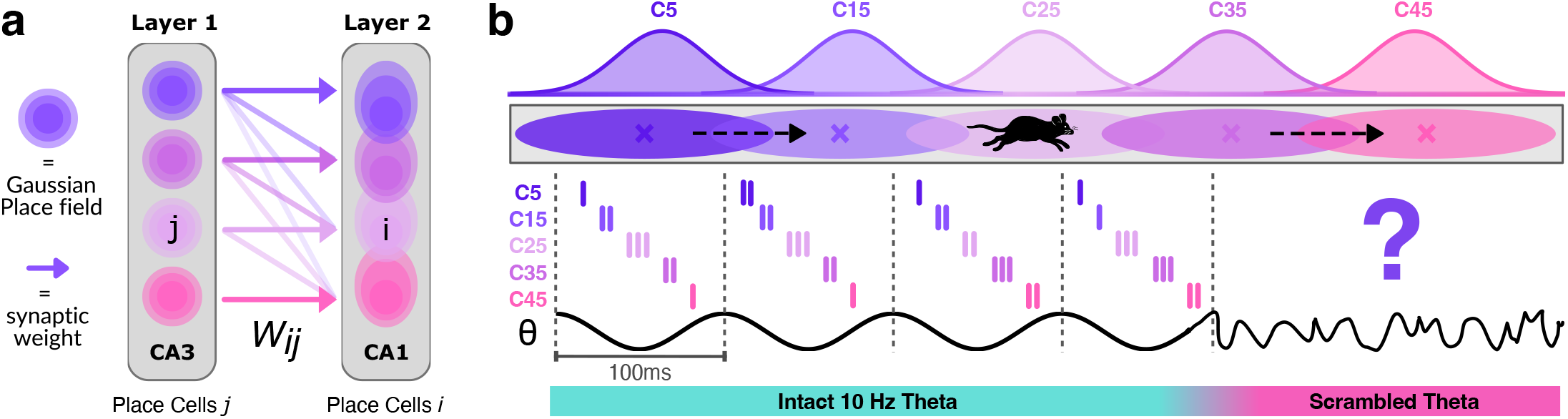
**a**, Schematic of the spiking neural network architecture, illustrating two layers of CA3 and CA1 place cells respectively and learned synaptic weights between them. **b**, Schematic of place cell activity forming theta sequences during intact oscillations while individual place cells encode the location of an animal on a unidirectional track running left to right.

The firing rate of any CA3 cell *j* at agent position *x* at time *t* is given by:

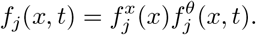

Activity in downstream CA1 place cells is directly driven by CA3 cell activity. Hence, phase precession and theta sequences are effectively inherited by CA1 cells. The firing rate of both CA3 and CA1 cells is transformed into spike trains through an inhomogeneous Poisson process, and the resulting spikes drive learning in the synaptic weight matrix *W*_*ij*_ between CA3 and CA1 cell populations according to an STDP learning rule (see Methods and George *et al*. [32] for more details). This weight matrix encompasses the predictive spatial map, as it encodes environmental transitions between states encoded by place cells and the expected occupancy of them discounted into the future. To evaluate the role of theta rhythms in learning and memory retrieval, we implemented three distinct manipulations of the oscillation. The control condition is an intact 10 Hz oscillation given by:

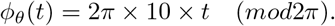

A second manipulation implements frequency scrambling as performed experimentally in in vivo studies using optogenetics [30]. This was implemented as circular Brownian motion with drift defined as:

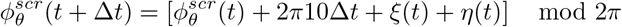

where the drift term 2*π*10Δ*t* describes an intact 10 Hz oscillation and Brownian noise is added through two noise terms, *ξ*(*t*) and *η*(*t*) at each time step. These parameters allowed us to control the amount of frequency scrambling from 0 to 1 (Fig. 2**a**) and effectively abolished theta power for the highest values of scrambling (Fig. 2**b**). Since the relationship between the amount of scrambling and *R*^2^ values were non-linear (Fig. 2**b-c**), later analyses implement a log-scale when reporting results. Finally, a third condition entirely removes theta oscillations, leaving only the pure spatial rate codes.

**Figure 2:**
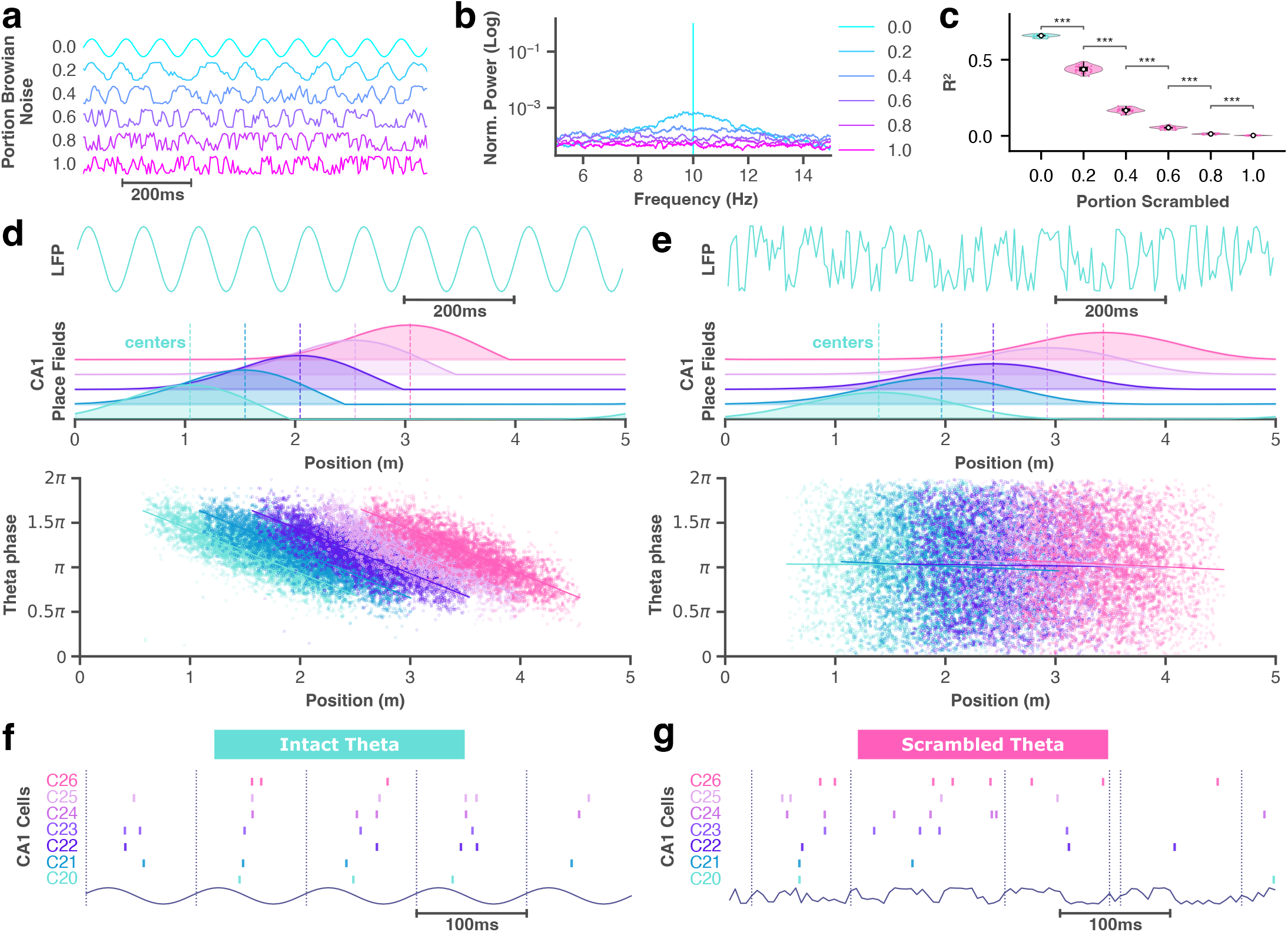
Scrambling theta oscillations disrupts theta sequences, phase precession, and learned representations. **a**, Example theta oscillations with varying amounts of additive noise. 0 refers to a pure theta sine wave and 1 to pure Brownian noise. **b**, Corresponding fast Fourier transform power spectra. Spectra of scrambled conditions are normalized to the peak of the intact theta signal for visualization purposes. **c**, Distribution of *R*^2^ values of fitted regression lines for selected cells across different degrees of theta frequency scrambling. Higher *R*^2^ values correspond to stronger theta phase-locking, which is linked to higher theta sequenceness. Data represent median (white dot) ± upper and lower quartile, ns – not significant, *p<0.05, **<0.01, ***<0.001. **d**, Reference oscillation (top), CA1 place fields with dotted lines indicating centers (middle) and spiking patterns of selected neurons across phases and positions on the 5 meter track (bottom) in the intact theta condition. **e**, Same for the pure scrambled theta condition. **f**, Example theta sequences during intact theta. **g**, Same for the scrambled theta condition.

Next, we evaluated the impact of theta frequency scrambling on theta sequences and spike timing. To this end, we simulated an agent with 50 evenly spaced CA3 place cells on a 5 meter linear track where the right end loops back into the left end, effectively forming a closed infinite loop. The agent explored the maze for 30 min moving left to right at a constant velocity of 0.16 cm/sec. This simulation was run under both intact 10 Hz (Fig. 2**d**) and fully scrambled theta (i.e. *σ*_*ξ*_ = *σ*_*η*_ = 1.0; Fig. 2**e**). While intact theta generates stable phase precession, reflected in a clear relationship between oscillation phase and spike timing, with cells spiking at progressively earlier phases as the agent advances along the track (Fig. 2**d**), under frequency scrambling, that relationship degrades (Fig. 2**e**). To more specifically quantify how scrambling affects phase locking, which we express as the coefficient of determination (*R*^2^) of each cell’s phase-position regression line, we analyzed the *R*^2^ values for different degrees of frequency scrambling using a beta generalized linear mixed model with scrambling as a fixed effect, cell ID as a random intercept (accounting for the same cells being simulated under 6 levels of frequency scrambling), and dispersion being allowed to vary between levels (see methods). Higher *R*^2^ values correspond to stronger phase-locking and vice versa. Frequency scrambling had a significant effect on phase locking (*R*^2^) (Wald *χ*^2^(5) = 28, 773, *p <* .001). More specifically, model estimated mean *R*^2^ values decreased at every modeled step of increased scrambling from 0.66 (95%CI[0.65;0.66]) under intact theta to 0.44 (95% CI[0.43;0.45]), 0.17 (95% CI[0.16;0.17]), 0.053 (95% CI[0.050;0.056]), 0.013 (95% CI[0.012;0.015]) and 0.003 (95% CI[0.002;0.004]) at scrambling degrees 0.2 to 1.0 (Fig. 2**c**). Post-hoc pairwise comparisons (estimated marginal means, Tukey-adjusted) revealed significant differences in mean *R*^2^ values between every pair of modeled scrambling degrees (all *p*_adj_ *<* .001; Fig. 2**c**)). Furthermore, the decline was not uniform across degrees of scrambling, but largest at low to intermediate scrambling degrees (e.g. Δ*R*^2^ = 0.27 between 0.2 and 0.4) and became smaller at higher degrees of scrambling (Δ*R*^2^ = 0.01 between 0.8 and 1.0) as *R*^2^ approached its floor of zero (Fig. 2**c**), suggesting that larger changes might already happen at lower degrees of frequency scrambling. These results suggest that although firing rates remain temporally modulated, the lack of a coherent theta rhythm prevents phase-locking across agent positions and increasingly so as frequency scrambling increases. This disruption is further visible at the level of individual theta cycles, where theta sequences associated with 10 Hz oscillations (Fig. 2**f**) degrade during frequency scrambling (Fig. 2**g**). It should be noted that, regardless of the temporal disruption of cell activity, CA1 place fields, reflecting the rate code of place cells at any agent position on the track, can still be derived from the encoded successor representation and overall maintain their spatial selectivity (Fig. 2**d-e**), with CA1 place field centers learned under intact theta showing a slight backwards shift (Fig. 2**d**) compared to CA1 place fields learned under fully scrambled theta (Fig. 2**e**). Altogether, these results suggest that frequency scrambling disrupts the temporal organization of spikes, resulting in impaired phase precession and disorganized within-cycle sequences, while leaving broad locational information largely intact.

### Theta frequency scrambling impairs learning predictive spatial representations

Since theta oscillations are required for successful learning of predictive spatial maps [32] while leaving the exact role of theta rhythmicity undefined, we aimed to address this question using our frequency scrambling approach. To this end, we employed the same maze type, parameters and exploration time as outlined in the previous section, but additionally also ran simulations under no theta. During the exploration time, synaptic weights in the weight matrix *W*_*ij*_ between CA3 and CA1 cell populations were learned through STDP, encoding a successor representation (SR). Under intact theta, this learned predictive representation exhibits a visible backward skew opposite the direction of travel as indicated by a negative skewness value (skew = - 0.48). Here, skewness is expressed on a symmetric scale from -1.0 to 1.0, where 0 indicates perfect symmetry. Specifically, the backwards skew is reflected in an off-diagonal left shift of the diagonal positive weight band in the matrix (Fig. 3**a**). This asymmetry reflects learned successor representations [32, 33]. Crucially, this shift was less pronounced after learning under fully scrambled theta (skew = -0.26) (Fig. 3**b**) and practically abolished for the SR learned under no theta (skew= -0.02) (Fig. 3**c**). This trend is further apparent in the distribution of synaptic weights across states encoded in the SR (Fig. 3**d**), which shows a visible asymmetry and left shift of the peak of the distribution under intact theta, while no such shift is observed under scrambled or no theta leaving spatial representations broader and more symmetric. Furthermore, we observed a linear trend for skewness values across different degrees of scrambling, where increasing degrees of frequency scrambling were significantly associated with lower backward skewness of the SR (*R*^2^ = 0.94, *p* = 0.001). Additionally, similarity between weight matrices learned under scrambled and no theta (*R*^2^ = 0.83) was higher compared to the similarity between weight matrices learned under intact and scrambled theta (*R*^2^ = 0.39) as well as intact and no theta respectively (*R*^2^ = 0.21). While the lack of a predictive backwards skew under no theta compared to intact theta is a direct replication of findings reported by George *et al*. [32], our frequency scrambling paradigm further suggests that intact theta rhythmicity is a decisive factor in learning appropriate predictive representations.

**Figure 3:**
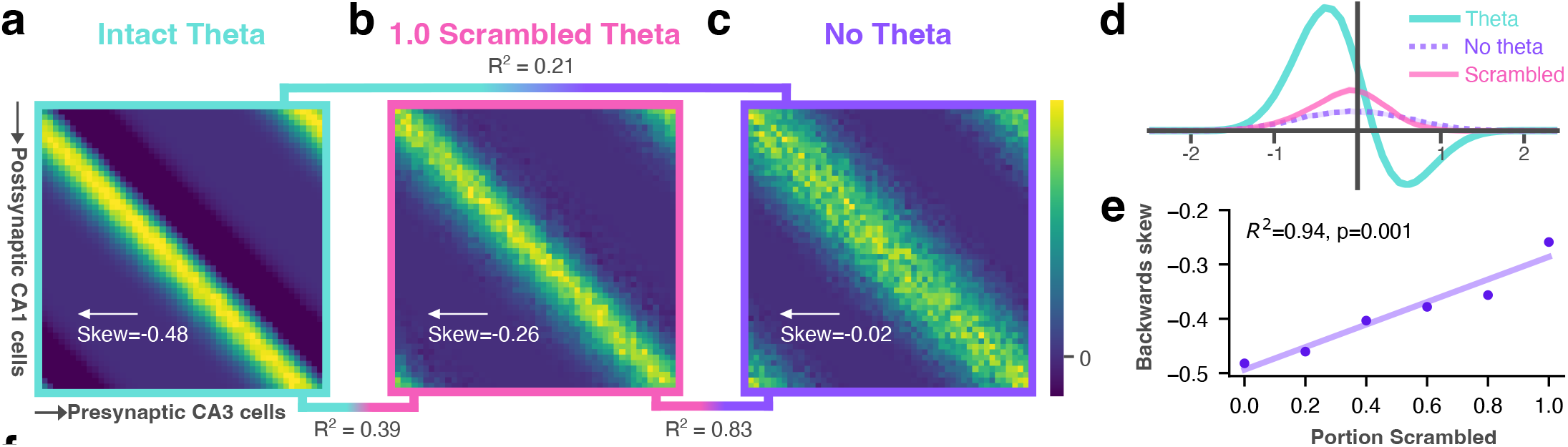
**a**, Synaptic weight matrix (SR) learned under intact theta. Presynaptic CA3 and post synaptic CA1 cells are ordered in rows and columns according to their position on the track. **b**, Same for maximally scrambled theta. **c**, Same for no theta. **d**, Synaptic weight distributions for all theta manipulations. Rows of the respective weight matrix were aligned on the diagonal and subsequently averaged to obtain the distribution. **e**, Skewness values obtained from the weight matrices across different degrees of scrambling. The trend between skewness and increasing scrambling is visualized by the regression line.

### Theta frequency scrambling impairs spatial memory retrieval

Following our findings on the role of theta in memory encoding, we next evaluated the role of theta sequences in memory retrieval. To this end, we trained an agent to navigate towards a reward in a T-maze (Fig. 4**a**). A set of 40 CA3 place cells were distributed along each arm and stem of the maze, resulting in a total population of 120 CA3 cells. To dissociate the role of theta in learning versus retrieval, the agent was pretrained for 30 min with an intact global 10 Hz theta oscillation to learn a predictive map *W*_*ij*_ that maximizes reward collection. During pretraining, the movement policy was the same as in the loop maze, meaning the agent moved in a straight line, but additionally took a 90°turn to the left or right depending on its arm choice at the T-maze junction. To control agent exploration and balance correct choices while allowing for the exploration of the entire maze, we programmed the agent to choose the correct arm with a fixed probability of 0.9, drawn independently on each trial from a uniform distribution. Consequently, the performance ceiling for the agent on the subsequent navigation task is also 90% accuracy. After pretraining, the weights in *W*_*ij*_ were frozen, separating learning from subsequent retrieval. During the following navigation task, the agent’s movement was controlled by downstream CA1 place cell spikes, reflecting the readout of the learned predictive map *W*_*ij*_. At each position of the agent, a direction vector was computed from spikes occurring in the late phase of theta cycles, reflecting possible future positions, that determines the direction of the agent’s next step (see Methods for more detail; Fig. 4**b**). Since theta sequences encode possible future locations, but not necessarily how they relate to rewards in the environment, an additional value function (Fig. 4**c-d**) is given to the agent, which assigns a value to locations encoded by a place cell based on the cells distance to the reward. Although in this task we manually choose how late phase spikes control behavior, which potentially induces some form of circularity, this does not invalidate the findings given that our main aim here is purely to decode hippocampal output to establish the effects of frequency scrambling. Recreating hippocampal activity, exposing it to frequency scrambling and decoding, and visualizing the subsequent readout provides a targeted test of this effect. Task simulations are run as episodes that time out after 90 seconds, unless the agent reaches the reward or chooses the wrong arm before that time limit is reached (see Fig. 4**e** for example trajectories of each case). A total of 50 episodes (5 seeds with 10 episodes each) are run for different theta manipulations, respectively. Specifically, we simulate task performance under an intact 10 Hz theta oscillation, as well as under different degrees of frequency scrambling. We ran simulations under evenly spaced values of scrambling ranging from 0 (intact theta) to 1.0 (fully scrambled). However, since no major differences were observed between higher values of scrambling (see Fig. S**1**), we spaced the degrees of scrambling logarithmically rather than linearly to sample more densely at lower values of scrambling and better visualize effects in that range (see methods for details). The log values were used in any results reported below.

**Figure 4:**
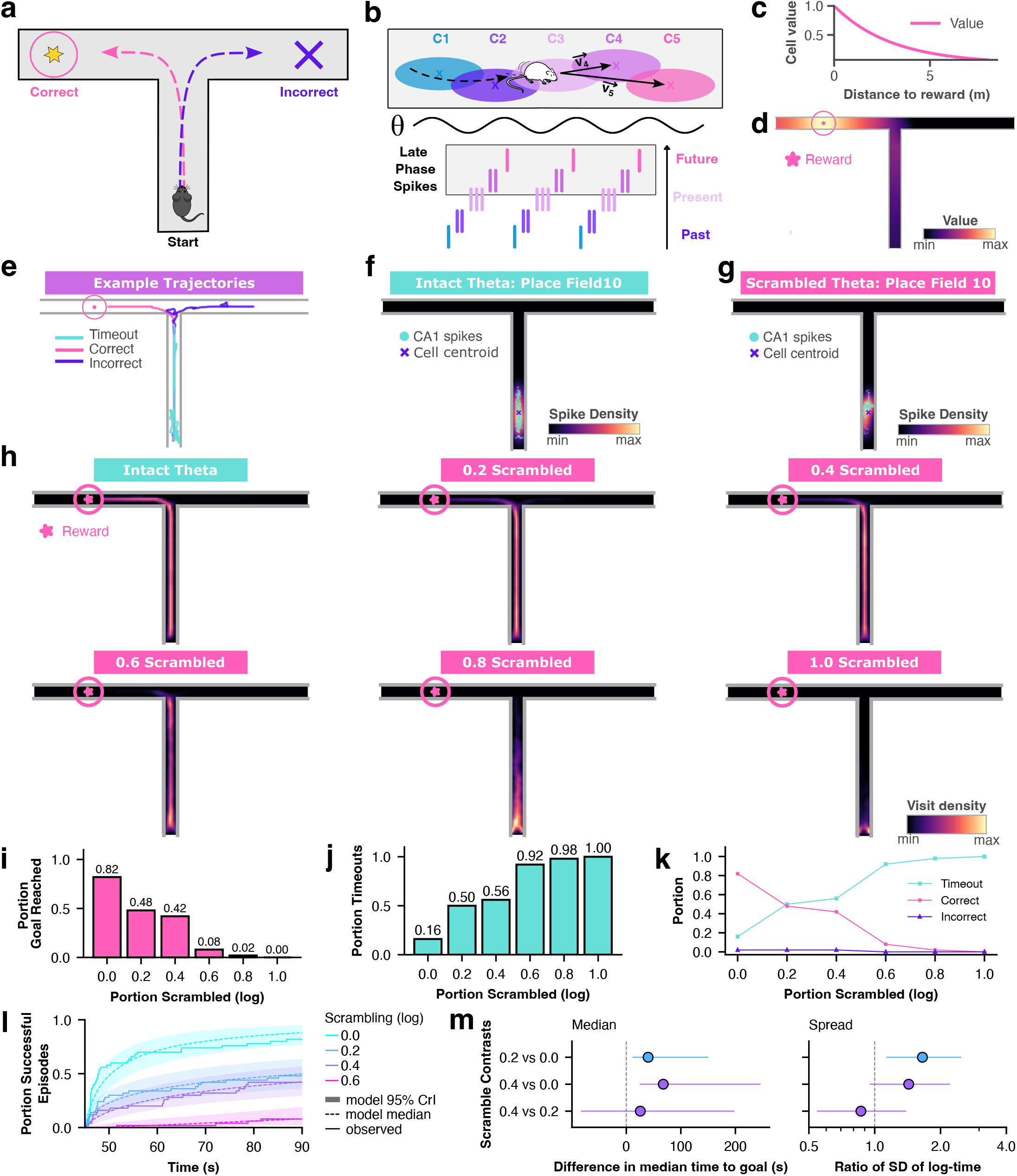
**a**, Memory-guided choice task. **b**, Schematic of movement policy: Direction vectors are decoded from late-phase spikes (indicated by the light gray rectangle), determining agent movement direction. **c**, Value function: Cell values as a function of distance to reward. **d**, Value function: cell values interpolated across maze locations. **e**, Example trajectories. **f**, Place field of a selected cell under intact theta reconstructed from CA1 spikes (cyan dots). **g**, Same for a place field under fully scrambled theta. **h**, Trajectory heatmaps across episodes and different degrees of frequency scrambling (log scale). **i**, Portion of correct arm choices across scrambling degrees. **j**, Same for the portion of timeouts. **k**, Performance metrics across conditions. **l**, Navigation efficiency: Cumulative portion of successful episodes over time. **m**, Model-estimated differences in median time to reward and ratios of SDs between conditions (95% CrI).

Given that hippocampal place fields remain intact despite theta frequency scrambling [30], CA1 place fields were reconstructed from episode-wise spike data to assess whether we could replicate this dissociation. During exploration under both an intact theta oscillation (Fig. 4**f**) and maximum scrambling (Fig. 4**g**) place fields remained intact, indicating that CA1 neurons retained their spatial selectivity also during substantial disruption of the theta rhythm. In contrast, increasing frequency scrambling considerably altered the agent’s movement and navigation behavior (Fig. 4**h**). Under an intact theta oscillation, the agent mostly traveled straight to the maze junction before choosing one of the two arms. However, with increasing frequency scrambling, trajectories became progressively less structured as well as less goal-directed, and exploration of the maze remained increasingly constrained to the vicinity of the starting position, suggesting impaired planning and goal-directed navigation (Fig. 4**h**). These behavioral changes were also reflected in task performance (Fig. 4**i-l**). Under intact theta, the agent successfully reached the reward in 82% of episodes across seeds, with the remaining episodes being either incorrect choices (2%) or timeouts (16%). Increased frequency scrambling progressively reduced the proportion of successful episodes (Fig. 4**i** 48%, 42%, 8%, 2% and 0% at scrambling strengths *s* = 0.2, 0.4, 0.6, 0.8 and 1, on the log scale), while increasing the number of unsuccessful trials, including timeouts (Fig. 4**j-k**).

To quantify navigation efficiency, we modeled the time to reach the reward using Bayesian censored log-normal regression, which accounts for episodes that ended at the 90 s time limit without reaching the reward (see methods for details). Navigation efficiency deteriorated quickly, even for low scrambling values (Fig. 4**l**). Under intact theta, the model estimated median time to reward was 50.3 s (95% CrI [47.9, 54.9]). At 0.2 scrambling this time increased by 40.0 s (95% CrI = [11.69; 150.38]), and at 0.4 scrambling by 67.8 s ([25.9, 245.8]), relative to intact theta (Fig. 4**m**). The difference between 0.2 and 0.4 was however insignificant (+25.7 s, [− 83.2, 198.1]), consistent with their overlapping curves in Fig. 4**l**. Additionally, episode completion times became also significantly more variable and inconsistent at scrambling of 0.2 compared to intact theta (1.65; 95% CrI [1.13, 2.49]) but not when comparing 0 and 0.4 or 0.2 and 0.4 (Fig. 4**m**). Since less than 50% of episodes were completed successfully for scrambling values ≥ 0.4, the estimated medians for these conditions lie beyond the 90 s episode time limit and are model-based extrapolations, explaining why the corresponding credible intervals are wide. Furthermore, scrambling values ≥ 0.8 were excluded from the contrasts given their lack or very limited number of successful episodes, but were retained in the analysis of the overall trend. Together the results show that frequency scrambling, and the associated disruption of theta sequences, impaired both the accuracy and efficiency of spatial navigation in the model, and that even low degrees of scrambling can result in significant deficits that become only more substantial as scrambling strength increases.

## Discussion

While the relevance of theta oscillations for learning and memory was long-known, our results provide a potential specific mechanism for why that is the case, by linking theta oscillations and more specifically theta sequences to learning and online retrieval. We show that frequency scrambling effectively disrupts both phase precession and theta sequences in CA1, resulting in impaired read-out of the encoded spatial information and degraded navigation performance, expressed in decreased movement stability and efficiency. Thus, the disruption of hippocampal spatiotemporal codes can compromise place-cell dependent navigation, even when the underlying spatial representations remain intact.

### Beyond place cells as substrates of memory

Hippocampal place cells were long thought to be a key neural substrate of spatial memory. Their ability to reliably encode specific locations in space made them the perfect cellular elements comprising the spatial cognitive map, with their representations having been linked to navigation performance, but also memory in a broader sense, given that space is a central element in both, episodic and semantic recollections of experience [3, 4, 34–36]. However, in line with previous work [30, 31], we show that memory retrieval can be impaired despite intact spatial representations encoded by place cells, challenging the idea that place cell physiology alone is sufficient to explain memory. Specifically, while the hippocampal ‘cognitive map’ has historically been seen as primarily ‘spatial’[3, 4], our findings support the idea that this view does not fully capture the process of memory encoding and retrieval, particularly in real-time, and that instead, an added temporal dimension plays a crucial role in the underlying mechanisms. This idea dates back to Tulving [37], who proposed that memories are embedded in both a spatial and a temporal context, and is supported by empirical and theoretical evidence showing that representations of spatial memories are actively organized into temporally ordered sequences and that the hippocampus contributes to remembering the order of events [38]. More broadly, this suggests that the hippocampus organizes representations in time to form associations between sequential events, with space providing the relevant contextual background for this process [39]. In this view, theta oscillations provide a plausible mechanism for orchestrating this temporal organization, and theta sequences, together with synaptic plasticity rules, may constitute the format through which events are bound in a directional associative way within a spatiotemporal context.

### Real-time memory retrieval during navigation

Another key implication of our findings is that efficient memory retrieval during navigation depends on temporally organized neural activity on the millisecond timescale. Although choices during navigation must often be made rapidly, place fields are typically computed from neural activity sampled over multiple minutes of behavior [30]. This suggests that spatial rate codes alone may provide useful information only over longer timescales, whereas theta sequences provide a compressed temporal online code that can efficiently represent candidate trajectories on the millisecond scale, and thereby support real time memory retrieval. This idea is supported by a recent study proposing theta sequences as a suitable neural substrate for online goal-directed planning [19]. Furthermore, another recent study used neurofeedback to reward spatially remote representations in a Y-maze to link hippocampal CA1 activity patterns occurring predominantly at the late theta phases to real-time memory retrieval and mental teleportation [40]. Finally, theta sequences have also been proposed to contribute to retrieval in hippocampus-prefrontal cortex networks during spatial tasks, where precise temporal coherence and phase-locked spiking to theta oscillations appear crucial [41]. Accordingly, when this temporal organization is disrupted, the spatial map may remain intact in a rate-based sense, but its behavioral readout becomes less effective for guiding contextually appropriate actions.

### Theta sequences as a temporal scaffold for retrieval

Overall, our results show that frequency scrambling of theta oscillations disrupts the temporal structure of place cell activity and, as a consequence, the continuous representation of sequential experience as the animal explores an environment. It has long been argued that this organization is crucial for episodic memory formation as it allows synaptic plasticity rules, such as STDP, to bind successive events together and encode the temporal and thus contextual relation between them [14, 39]. Our findings suggest that the same compression of consecutive states into sequences is also required for the retrieval and use of encoded spatial representations. When the rhythmic and predictable structure of theta oscillations is lost, their readout becomes less effective with likely consequences for downstream regions that must interpret and integrate hippocampal CA1 output. Hence, rather than viewing the hippocampus as a static spatial map, our results support a framework in which spatial information encoded by place cells must be reactivated in a temporally structured format to enable effective readout by downstream regions and behaviorally appropriate use. This interpretation also aligns with the predictive map hypothesis, which proposes that the hippocampus represents environmental transitions and consequently also future states [32, 33]. Recent work, as well as our results, show that theta oscillations are crucial for learning such predictive maps and that in their absence, the learned representation loses its predictive structure [32]. Together, these findings suggest that the temporal organization that supports learning of environmental transitions by binding sequential experience must also be reinstated during retrieval for the representation to guide navigation effectively.

### Relevance for in vivo systems

Questions about the role of the hippocampal formation in navigation, planning and memory are to this day largely addressed in vivo [19–21] and it is likely that these approaches will continue to be indispensable to advancing our understanding of the living brain. Yet, in vivo research is not free of limitations and challenges, especially when it comes to the controlled isolation of specific circuit dynamics or the impact of an intervention on targeted brain areas and functions. In vivo it is difficult to manipulate one factor, such as the timing of theta, while leaving everything else in the brain unchanged. Additionally, techniques are often invasive and require the use and sacrifice of animals. Computational approaches, such as the one proposed here, hold the potential to contribute to establishing the sufficiency of physiological principles while contributing to the refinement and reduction of animal experimentation, which aligns with current and future legislation around animal experimentation [42, 43]. Specifically, these approaches have the capacity to isolate and manipulate certain aspects of brain function in a highly controlled manner. In our model for example, the temporal organization of theta could be scrambled selectively and gradually, while task, place cell rate codes and readout were held fixed, enabling us to isolate its possible contribution to navigation in a way that is difficult to achieve in a living brain. While in vivo experiments will remain necessary, in silico approaches could suggest more targeted studies and reduce the need for animal studies overall, with some studies potentially being shifted to computational model entirely. The present study is an example of this approach. It tests a hypothesis derived from in vivo findings [30], isolates a mechanism that has been difficult to dissociate experimentally, and yields predictions for subsequent more targeted in vivo studies, such as the graded impairment of memory retrieval and loss of navigation success and efficiency with increasing disruption of theta rhythmicity.

### Limitations

Our model only captures a simplified and isolated neural navigation circuit, in which behavior is directly driven by CA1 output and does not explicitly incorporate broader entorhinal, cortical, sensory, or reward-related networks. As a result, it cannot capture the full distributed computations across circuits that support memory processes during navigation and decision-making. At the same time, this simplification is useful in that it specifically isolates the role of theta-dependent temporal organization of place cell activity and allows us to assess how its disorganization affects the retrieval of hippocampal spatial representations. In live rodents, memory-guided navigation depends on the coordinated interactions between the hippocampus and downstream regions, specifically the medial prefrontal cortex (mPFC), which is thought to integrate hippocampal information in decision-making [44]. Specifically, theta synchrony and phase-locking of spikes between both regions have been linked to memory-guided choices, prediction of goal locations [45] and shaping as well as continuously updating of cortical spatial representations [44, 46, 47]. Accordingly, temporally disorganized CA1 output in our model may impair not only local sequence structure, but also the transfer of sequentially ordered spatial information to cortical networks, involved in evaluating candidate trajectories and integrating them with task demands or reward information. A similar caveat applies to the medial entorhinal cortex (mEC) which contains grid cells that are thought to provide a spatial metric for path integration and distance estimation [48, 49] and whose activity depends on intact theta organization [50]. While place fields remain largely intact under theta frequency scrambling [30], grid cell periodicity is strongly disrupted by medial septal theta manipulations [50]. This suggests that future research should further investigate their contribution to spatial memory encoding and retrieval during navigation. Furthermore, our model simplifies the relation between CA1 output and reward-guided decision making. We represent value with a predefined function that is typically biologically learned through experience, involving regions like the ventral striatum and other reward related circuits [51, 52]. Nevertheless, this simplification is consistent with the successor representation framework, which separates learning of environmental transitions from reward computations [53]. Likewise, ramp cells in the ventral striatum may integrate temporally structured hippocampal output to reflect proximity to rewards, matching the segregation between theta sequences and the value function in our model [52]. It remains up to future research to further examine the exact mechanism underlying the integration of reward signals with spatiotemporal CA1 codes. Finally, previous studies using optogenetic silencing of theta oscillations have shown that the behavioral consequences of theta disruptions depend on the task period. Specifically, Gemzik *et al*. [29] and Koenig *et al*. [50] showed that while theta suppression during the delay period, relevant for working memory, impairs memory maintenance, theta silencing during the retrieval period had somewhat weaker effects on memory-guided task performance in spatial navigation. By contrast, frequency scrambling specifically disrupts theta rhythmicity while preserving general theta activity, making it a more targeted manipulation of the theta sequence structure. This could explain why theta silencing and theta frequency scrambling may differently affect memory retrieval and produce disparate behavioral consequences, despite both manipulating theta oscillations. In conclusion, our results indicate that theta sequences provide a spatiotemporal code that is critical for the effective online retrieval and use of spatial representations during navigation. Although theta sequences are currently a major focus in hippocampal research [19–21, 54], their precise role in goal-directed navigation, online planning and memory process remains to be fully defined.

## Methods

### Summary of the model

We adopted the spiking neural network model developed George *et al*. [32] and extended it for the present task, while keeping the core architecture unchanged. The model consists of a layer of *N* CA3 basis feature cells *f*_*j*_(*x, t*) for *j* ∈ {1…*N*} that encode the current position *x*(*t*) of an agent exploring a maze, and a downstream layer of CA1 successor feature cells *ψ*_*i*_(*x, t*) whose activity is driven by the spiking of the upstream CA3 cells. Each CA3 basis feature cell has a spatial receptive field defined by a thresholded Gaussian distribution with mean *x*_*j*_ and standard deviation *σ* = 1*m*, where *x*_*j*_ denotes the center of the receptive field. The position of the agent within the receptive field is encoded by the spatial firing rate of the cell, which reaches its maximum rate *F* at the center *x*_*j*_ of the field and is 0 outside of the field:

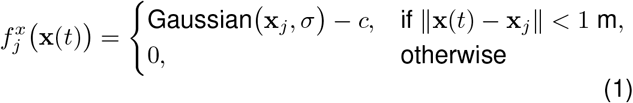

The peak firing rate was set at 10 Hz which is around experimental values reported by Mizuseki *et al*. [55]. The exact firing rate of a CA3 basis feature *j* at any location *x*(*t*) of the agent, according to the pure spatial rate code, is given by:

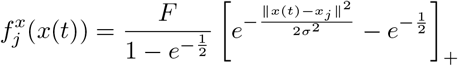

Phase precession, as implemented in the model, suppresses the firing rate for all but the preferred phase of a CA3 cell in the theta cycle. This preferred phase shifts gradually backward as the agent traverses the cell’s receptive field, being late as the agent enters the field and early as the agent is about to leave the field. This effectively gives rise to phase precession at the individual cell level and theta sequences at the population level. Mathematically this is implemented through a phase precession factor by which the spatial firing rate is multiplied (explained in further detail below). The resulting theta modulated firing rate of both populations, (*f*_*j*_(*x, t*)) and (*ψ*_*i*_(*x, t*)), is transformed into spike trains through an inhomogeneous Poisson process, and these spikes drive online learning in the synaptic weight matrix between the CA3 and CA1 cell populations according to an asymmetric Hebbian STDP learning rule. In simple terms, this rule states that if a presynaptic CA3 cell *j* spikes shortly before a postsynaptic CA1 cell *i*, the weight *W*_*ij*_ between them is strengthened (potentiated), whereas when the postsynaptic cell spikes before the presynaptic cell, the weight is weakened (depressed). The model implements this by using eligibility traces for presynaptic basis features 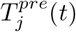and postsynaptic successor features 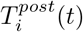. These traces increase with each respective spike from a cell *j* or *i* and decay otherwise according to a plasticity window with decay times of *τ* ^*pre*^ = 20 ms for presynaptic cells and *τ* ^*post*^ = 40 ms for postsynaptic cells. The decay times were adopted from George *et al*. [32] and stem from experimental data from Bush *et al*. [56]. Synaptic weights were updated over the exploration period of the agent in the maze, using the accumulated spike timing differences and using default parameters reported in George *et al*. [32], unless specified otherwise:

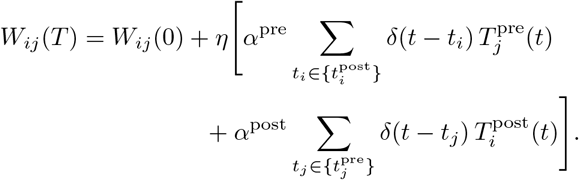

### Implementation of phase precession and theta sequences

The model implements phase precession through a phase precession factor by which the pure spatial rate code 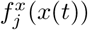 is multiplied:

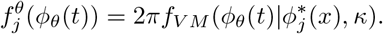

where *f*_*V M*_ (*x*|*µ, κ*) is the circular Von Mises distribution of a CA3 cell *j* at the current theta phase *x* ∈ (0, 2*π*), with mean 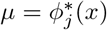, denoting the preferred phase of that cell at the agent’s current position and spread parameter *κ* = 5, which controls the degree of phase locking, that is, the margin around the preferred phase within which spikes occur. Lower kappa values are associated with a wider spread, leaving more leeway for spikes to occur at phases further from the preferred phase. In contrast, higher kappa values result in a more concentrated Von Mises distribution, with spikes being increasingly constrained to occur at or closer around the preferred phase. The phase precession factor is only large when the phase of the global theta oscillation *ϕ*_*θ*_(*t*) at time *t* is close to the preferred phase 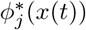 of a cell *j* at position *x* of an agent at time *t*. The final firing rate of a CA3 basis feature *j* is the product of the spatial rate code based firing rate and the phase precession factor:

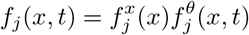

This effectively leads to suppression of the firing rate for all but the preferred phase with a certain margin determined by *κ*. In the model, theta sequences arise directly from phase precession: As the agent traverses a cell’s receptive field, spikes shift to progressively earlier phases within successive theta cycles, effectively generating sequences at the population level that represent past, present, and potential upcoming locations along the agent’s trajectory. Sequence clarity is controlled by the Von Mises spread parameter *κ*, with higher values corresponding to stronger phase locking and sharper sequences, whereas lower values allow for greater deviation from the preferred phase and thus leave sequences more diffuse. Since CA3 cell activity directly drives activity in the CA1 cell population, phase precession and theta sequences are inherited and thus present in both layers of the model.

### Implementation of theta manipulations

The model comprises three different manipulations of its global theta oscillation: Intact theta at a specified frequency (1), no theta (2), and a frequency scrambled oscillation (3).

The default intact global theta oscillation is defined as a sine wave with phase *ϕ*_*θ*_ at time *t* given by

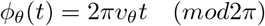

where the frequency *v*_*θ*_ of the oscillation defaults to 10 Hz, and *ϕ*_*θ*_(*t*) ∈ [0, 2*π*].

Under the “no theta” manipulation, no temporal modulation of the spatial rate code 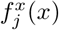 takes place, meaning that the spiking activity of the CA3 basis feature cells depends entirely on the firing rate defined by the Gaussian place field and the agent’s position *x* within that field. Consequently, synaptic weights in the SR are learned via the STDP learning rule, using the pure spatial rate code given by

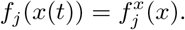

The frequency scrambled theta oscillation was designed to generate a signal that mirrors the effects of optogenetic frequency scrambling on hippocampal oscillations, as observed experimentally in in vivo studies such as Etter *et al*. [30]. To this end, phase scrambling of the intact 10 Hz theta oscillation *ϕ*_*θ*_(*t*) was modeled as a discrete-time circular Brownian motion with drift. At each simulation time step Δ*t*, the phase was advanced and stochastically perturbed using the following equation:

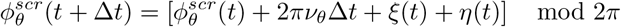

where 2*πν*_*θ*_Δ*t* denotes the drift term with oscillation frequency *ν*_*θ*_ and time step Δ*t*. Stochastic noise at each time step was introduced by two independent and identically distributed zero-mean Gaussian distributions *ξ*(*t*) ∼ *N* (0, *σ*_*ξ*_) and *η*(*t*) ∼ *N* (0, *σ*_*η*_), with *σ*_*ξ*_ = *σ*_*η*_ ∈ [0, 1]. Since *ξ*(*t*) and *η*(*t*) are independent and identically distributed at each time step Δ*t* , their combination essentially results in a single phase perturbation step that is described by a total variance of 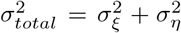 . Together, the two noise parameters control the degree of frequency scrambling. Consequently, in the absence of noise (when 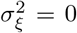 and 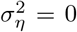) the equation simplifies to the intact, unscrambled theta rhythm, which is completely defined by the drift term. Finally, the modulo 2*π* wraps the phase around the unit circle [0, 2*π*], ensuring that frequency scrambling is compatible with the existing model architecture and temporal modulation of the spatial rate code remains possible. This implementation keeps the phase precessing characteristics of the modeled CA3 place cells segregated from the process that drives the oscillation and its manipulation and thereby mirrors the known septohippocampal organization and dissociation between drivers of the theta rhythm and spatial codes.

### Value function and movement policy

During the T-maze navigation task, the agent’s movement was directly controlled by the spiking activity of the CA1 successor feature cells. More specifically, we formulated a movement policy that uses the late phase spikes of recent theta cycles to decode a direction vector at each time point *t* that determines the direction of the next step the agent takes. The rationale for this approach is based on research suggesting that theta sequences are used for planning of behavior and movement trajectories [18, 22, 24] and that late phase spikes represent possible future locations of the agent and as such indicate prospective trajectories. Although not all planned behavior is translated into actual behavior, this approach provides a simple and straightforward way of visualizing planned behavior and the agent’s ability to use information encoded in CA1 theta sequences to navigate the maze. In the model, CA1 spikes are the read out of the learned SR, that is the weight matrix *W*_*ij*_ between the two model layers. The SR encodes environmental transition and stores expected occupancy of future states independently of rewards, meaning it tells the agent where it is likely to go, given its current location, but not where it is worthwhile to go in terms of rewards [53]. The strength of the SR is that it specifically separates the computation of the predictive map from the reward vector [53]. However, translated back to the CA1 read out, spiking activity might hence only tell the agent where it is likely to go next based on what the agent has learned, but not where it should go to obtain a reward. To account for this, we manually defined a value function that assigns a value *V*_*i*_ ∈ [0, 1] to each cell *i* that decays exponentially with distance to the reward and provides the agent with a relevant environmental reward structure. The value for any cell is given by:

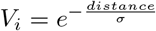

where *σ* = 3 determines the steepness of the decay and was chosen to ensure that cells in the stem, close to the starting point of the agent, have values *>* 0. To provide the agent with the information needed to make the correct arm choice (left arm), values of right arm cells (incorrect choice) were manually set to zero. The cell value is factored in when computing the direction vector at each time step. Mathematically, at any current position *x*(*t*) of the agent, we extract the late phase spikes from the last two theta cycles. To specify late phase spikes, we use a cut-off value of ≥ 1.5*π*, with any spike occurring at a phase ≥ 1.5*π* considered late. This value was chosen to ensure that most of the extracted spikes reflect future and not current or past locations. Spikes from the last two cycles are used to increase decodability of direction vectors from late phase spikes. Subsequently, a direction vector from the current agent position *x*(*t*) to the cell center *x*_*i*_ of the respective late-phase spiking CA1 cell *i* is computed:

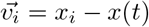

where *x*_*i*_ denotes the center of the *i*^*th*^ successor feature cell. Vectors are normalized to account for variation in distances between current location *x*(*t*) and the cell centers of late-phase spiking cells and ensure that vectors have equal magnitude:

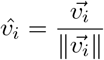

Resulting individual unit vectors are subsequently scaled by the spike count *s*_*i*_ of the successor feature cell *i* to which they point, accounting for the fact that greater cell activity, reflected by a higher spike count, should indicate a stronger direction signal. Additionally, vectors are weighted by a value term derived form the value function:

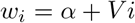

where *V*_*i*_ denotes the cell value and *α* = 1 is a constant that prevents the direction vector from being zeroed after multiplying when *V*_*i*_ = 0. This effectively ensures that in all cases *w*_*i*_ ≥ 1 and thus 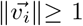. The complete individual vector from the agents current position *x*(*t*) to the cell center *x*_*i*_ of an active late-phase spiking CA1 cell, is given by

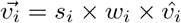

where, as mentioned above, *s*_*i*_ denotes the spike count of the *i*^*th*^ CA1 cell, *w*_*i*_ is the value term and 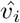 is the normalized initial individual vector. The final population direction vector 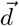 that drives the agent’s movement is the normalized sum of all individual vectors derived from the recent theta cycles, which is expressed by

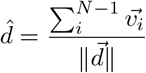

Individual vectors pointing at active cells closer to the reward or those with a higher spike count, have a larger influence on the direction of the population vector and consequently help guide the agent towards the reward. Since the magnitude of the population vector is always 1, the speed of the agent remains constant. In situations where the agent gets close to a wall or where a decoded population vector would lead to collision with a wall, the direction of the prospective step is changed parallel to the wall to bias the agent to follow walls, which mirrors thigmotaxis, a behavior observed in real mice and rats. This implementation has been adopted from the original model architecture as defined George *et al*. [32]. To make the overall movement trajectory more realistic, we furthermore set a maximum turn angle of 50 degrees to prevent extreme turns from one step to the next. When the angle between a direction vector at *t* − 1 (previous step) and *t* (prospective step) exceeded 50 degrees, the direction of the prospective step was adjusted to differ a maximum of 50 degrees from the direction of the previous step. Since the size of each movement step is very small, this however does not significantly restrict the agent’s movement freedom or its ability to find or not find the reward.

### Maze, task setup and simulation details

Simulations used one of two maze types: a loop maze or a T-maze. The loop maze is 5 meters long and structurally identical to a loop, with its right end looping back into the left end. 50 Gaussian CA3 basis feature cells with a radius of 1 m were evenly placed along the midline of the track. Due to the loop geometry, receptive fields close to the right end of the track extend into the left end and vice versa. Agents explored the maze for a total of 30 minutes according to a unidirectional left-right movement policy under which the agent moved around the maze in one direction at a constant speed of 16 ms/sec. The T-maze consists of a stem and two arms of 5 meters each. 40 Gaussian CA1 basis feature cells with a 1 m radius are placed in each of the arms and the stem with their centers evenly aligned along the midline of the maze. A reward was placed in the middle of the left arm 3 m from the maze junction with a radius of 0.5 m. During pretraining, the agent explored the maze for a total of 30 min according to a similar movement policy as in the loop, starting at the bottom of the stem and running up to the maze junction in a straight line. At the junction, the agent randomly chooses the left or the right arm with 90 to 10% probability respectively, effectively encoding the correct choice in the learned SR with some variation. Depending on the choice, the agent takes a 90 degree turn to either the left or the right and subsequently continues in a straight line until it reaches the reward or has traveled the same distance into the incorrect (right) arm. Upon reaching either of these points the agent is teleported back to the stem of the maze and the process is repeated until the training time is up. Importantly, during training, as well as during the subsequent task, the reward has no feedback effect on learning or performance and simply serves as a counter to quantify and keep track of the agent’s performance. Pretraining is completed under an intact 10 Hz theta oscillation. Once the agent has learned a predictive spatial map (SR), it performs the task using the late-phase spike based movement policy described in the previous section above. The task is performed under both an intact theta oscillation and an increasingly scrambled oscillation. Two sets of scrambling values were used for simulations. The first set was evenly and linearly spaced with *x* ∈ {0, 0.2, 0.4, 0.6, 0.8, 1.0}. The second set was logarithmically spaced and obtained by applying an exponential transformation

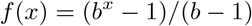

with *b* = 10 to each value *x* in the linearly spaced set, which puts values on a log_10_(1 + 9*s*) scale and yields *x* ≈ {0, 0.065, 0.168, 0.331, 0.590, 1.0}. This effectively places more values at lower scrambling, allowing for better differentiation of agent behavior in that range, while keeping minimum and maximum values at 0 and 1. During the task, the weights in the SR are frozen and no further learning takes place. We ran the task on 5 seeds with 10 episodes, resulting in a total of 50 episodes. For each episode, we record whether the agent reached the reward, chose the incorrect arm, or whether the episode timed out. In case of a correct choice, we also record the time needed to reach the reward. An episode timed out after 90 seconds, which is approximately twice the time the agent would need to attain the reward in a near perfect run. Data across episodes and seeds are pooled to obtain overall performance metrics.

### Statistical analyses

To verify our implementation of frequency scrambling, we plotted a trace of the oscillation for different degrees of scrambling (Fig. 2**a**) and conducted an analysis of their power spectra (Fig. 2**b**). Spectra were computed from 10 min of simulated theta rhythm for different scramble strengths and the resulting phase traces *ϕ*(*t*) were converted to an oscillatory signal *s*(*t*) = sin(*ϕ*(*t*)) to obtain a continuous time series signal suitable for spectral analysis. Power spectra were then computed using a one-sided Fast-Fourier Transform (FFT), which was applied to the mean-centered signal after Hann-windowing to reduce spectral leakage. For visualization, scrambled spectra were smoothed using a 0.2 Hz moving average, whereas the intact theta condition was unsmoothed to preserve the narrow spectral peak at 10 Hz. Furthermore, all spectra were normalized to the peak of the unscrambled condition to facilitate comparison across all conditions. To assess the effect of theta frequency scrambling on phase locking, we first decided to quantify phase-locking as the coefficient of determination (i.e. *R*^2^) of a regression line fitted to each cell’s spikes arranged in the phase-position plot in each scrambling condition. Due to the structure of the loop maze, the 30 cells around the middle of the loop (*x* = 2.5 *m*) were included in this analysis, since spikes of cells closer to one end reappear at the other end, resulting in skewed regression lines and inaccurate *R*^2^ values that would bias the results. Since the same 30 cells were simulated under all 6 levels of frequency scrambling and since values are bounded with *R*^2^ ∈ [0; 1], differences in *R*^2^ values were analyzed using a beta generalized linear mixed model (logit link) with condition as a fixed effect and and random intercept for cell, to account for the repeated measures structure. Additionally, dispersion was allowed to vary between conditions. The effect of frequency scrambling was tested with a Type III Wald *χ*^2^ test (analysis of deviance). Post-hoc contrasts were computed from estimated marginal means on the logit scale and correct for multiple comparisons using Tukey correction. Model fit was assessed with simulation based residual diagnostics (DHARMa). This analysis was completed in R using glmmTMB, car and emmeans. Navigation efficiency, operationalized as time to reward, was analyzed using a Bayesian censored log normal regression. Timed-out episodes, as well as the three wrong-arm choices, were treated as “duration longer than observed”, instead of being excluded, so that all episodes contribute information and conclusions are not limited to successes. Excess time, i.e. duration minus the minimum feasible time to reach the reward (fixed at 45 s based on the fastest observed success) were modeled as log-normal, with both mean and standard deviation of log-duration depending on scramble strength to account for unequal variances between conditions. A random intercept for seed was included. We fitted two models: A first model treating scrambling strength as a continuous predictor (inlcuding all six conditions) to assess the overall effect of frequency scrambling on navigation efficiency and a second model, treating scrambling conditions as categorical to examine differences between groups for conditions with at least some successes. Analyses were run using scrambling values from the logarithmically spaced set. Models were fitted with Bambi/PyMC (NUTS, 4 chains × 2000] post-warm-up draws, 2000 warm-up) using Bambi’s default weakly informative priors. In the continuous model, both the intercept (SD=3.6) and slope (SD=7.2) had zero-centered normal priors. In the categorical model, the condition-specific log-median excess time had zero-centered normal priors (all SD= 5.77). The coefficients of the model for the log SD of log-duration had *N* (0, 1) priors, and the standard deviation of the seed intercepts had a half-normal prior (scale 3.6 and 3.8) in the continuous and categorical and categorical models respectively. Convergence was checked through 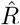 and divergences, and fit by comparing observed versus model-predicted success proportions and cumulative time-to-reward curves. We report posterior medians with 95% credible intervals (CrI) as well as contrasts reflecting differences in model-estimated median time to reward.

## Acknowledgments

GE is supported by a Proefdiervrij Pilot Grant. We thank Suzanne van der Veldt and Robbert Havekes for constructive discussions.

