## Supplemental Figure 1 for "Theta sequences support real-time spatial memory retrieval"

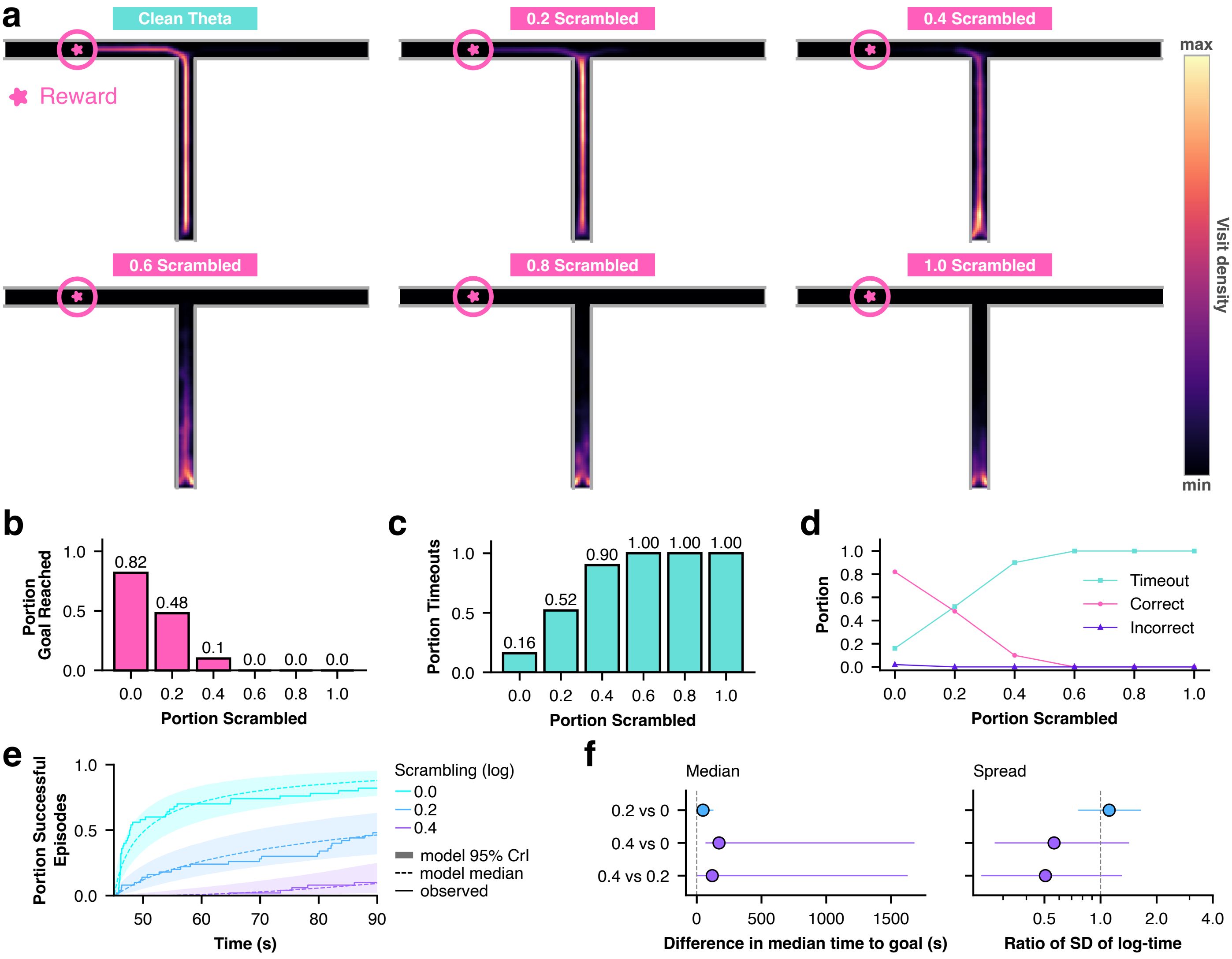


Figure S1: **a**, Trajectory heatmaps across episodes and different degrees of frequency scrambling values from the evenly spaced linear scale.**b**, Portion of correct arm choices across degrees of frequency scrambling. **c**, Same for the portion of timeouts. **d**, Performance metrics across conditions. **e**, Navigation efficiency: Cumulative portion of successful episodes over time for episodes with successes. **f**, Model-estimated differences in median time to reward and ratios of SDs between conditions (95% CrI). Credible intervals for values $\geq 0.2$ are strongly extrapolated (indicated by wide CrIs) given that the number of successes across episodes and seeds does not reach 50%.
